# Food touch limits lifespan through bioamine and neuroendocrine signaling

**DOI:** 10.1101/2024.04.19.590228

**Authors:** Elizabeth S. Kitto, Safa Beydoun, Scott F. Leiser

## Abstract

In multicellular organisms, sensory perception affects many aspects of behavior and physiology. Sensory cues are frequently perceived by the nervous system, which in turn coordinates systemic changes that can modulate health. Here we find that the sense of touch interacts with nutritional state to modulate lifespan in *C. elegans*. Worms subjected to dietary restriction are shorter-lived when they perceive tactile stimuli that mimic bacterial food and protective soil. Touch modulation of dietary restriction requires primary mechanoreceptors, the neurotransmitters dopamine and tyramine/adrenaline, and the neuropeptides insulin and GnRH. Ultimately, the touch circuit regulates the longevity effectors DAF-2/IGF1R and FMO-2/FMO5. These results establish a physiological touch circuit and connect neural reward pathways to the growth and reproductive axes. Finding that gentle touch can modulate longevity suggests a role for physical comfort in healthspan and lifespan.

## Main Text

In order to survive and thrive in a changing environment, all organisms must accurately perceive and respond to a wide range of sensory cues. Sight, touch, smell, taste, and sound inform animals of stressors, nutrient availability, and the presence of other organisms. Animals integrate this constant stream of information to respond appropriately to their environment: they must modify their behavior and metabolism to exploit favorable conditions while surviving or fleeing harsh conditions. In recent years, the field of geroscience has found that environmental stressors like dietary restriction (DR) ^1^, temperature stress ^2^, and hypoxia ^3^ can extend lifespan and healthspan . There is great interest in leveraging these longevity-promoting conditions to improve human health. The most well-studied of these interventions is DR, defined as a reduction in nutrient intake that does not cause malnutrition. DR extends healthspan and lifespan across taxa ^1^ and there is evidence it can benefit human health ^4^.

By genetically manipulating sensory perception, scientists have been able to decouple the perception of a stressor from exposure to the stressor itself. Surprisingly, perceiving a stressful environment can extend an organism’s lifespan, even under favorable conditions ^5^. On the other hand, perceiving attractive smells ^6-8^ and tastes ^9, 10^ decreases the health benefits of DR. Although previous work has shown that smell and taste modulate lifespan, other senses can also indicate the quality of the environment. For example, the sights and textures of food can signal abundant nutrition. Tactile cues can also indicate whether an environment is safe or dangerous, leading to changes in health and lifespan. For example, repeated tapping of the plate *Caenorhabditis elegans* (*C. elegans*) are cultured on decreases their lifespan under normal fed conditions through activation of a fight-or-flight response ^11^. But although organisms experience rich mechanosensory stimulation from food and shelter in their natural environment, the role of gentle mechanosensation in longevity is largely unexplored.

In this work, we utilize inedible food-like beads ^12^ to explore the role of touch in DR-mediated longevity in *C. elegans*. We find that touch modulates DR through a pathway that is distinct from smell, taste, and mechanosensory-driven behavioral circuits. These findings suggest a role for comforting and/or food-like touch in modulating lifespan. Specifically, our results implicate the neurotransmitters dopamine and tyramine/adrenaline as well as the hormones insulin and Gonadotropin-Releasing Hormone (GnRH) in touch modification of aging. These neural signals ultimately communicate touch information to the intestine, where the longevity effector FMO-2 modulates aging. The involvement of dopamine and tyramine/adrenaline suggests an interaction between motivational state and longevity. Moreover, these two neuroendocrine signals suggest a central role for the nervous system in balancing lifespan with growth and reproduction under stressful conditions via insulin and GnRH signaling, respectively.

### Mechanosensation attenuates DR-mediated longevity in an *fmo-2* dependent manner

In order to investigate the role of gentle touch on DR-mediated longevity, we sought an experimental condition that creates a tactile environment attractive to *C. elegans* like food and soil but lacking in other sensory cues. To separate the effects of touch perception from other nutrient cues, we utilized non-nutritive Sephadex beads to mimic the touch of bacterial food (**Fig. 1A**) ^12^. These beads are too large for *C. elegans* to consume (**Fig. S1A-C**), but they form a lawn-like texture similar to a real bacterial lawn. By combining the Sephadex “fake lawn” with fed and DR conditions (**Fig. 1A-B**), we were able to examine the effects of touch perception both with and without the smells and tastes of food. To confirm that this fake lawn is attractive to *C*. *elegans*, we measured lawn occupancy on fed and DR conditions with and without the fake lawn over a period of 24 hours. We found that *C. elegans* exhibit equal occupancy on the fed and fed + touch conditions but prefer to spend time in the center region of the plate under DR + touch relative to DR alone (**Fig. 1C**). This suggests that the fake lawn is attractive to *C. elegans* in a food availability-dependent manner and meets our criterion for a tactile mimetic of food and soil.

**Fig. 1.**
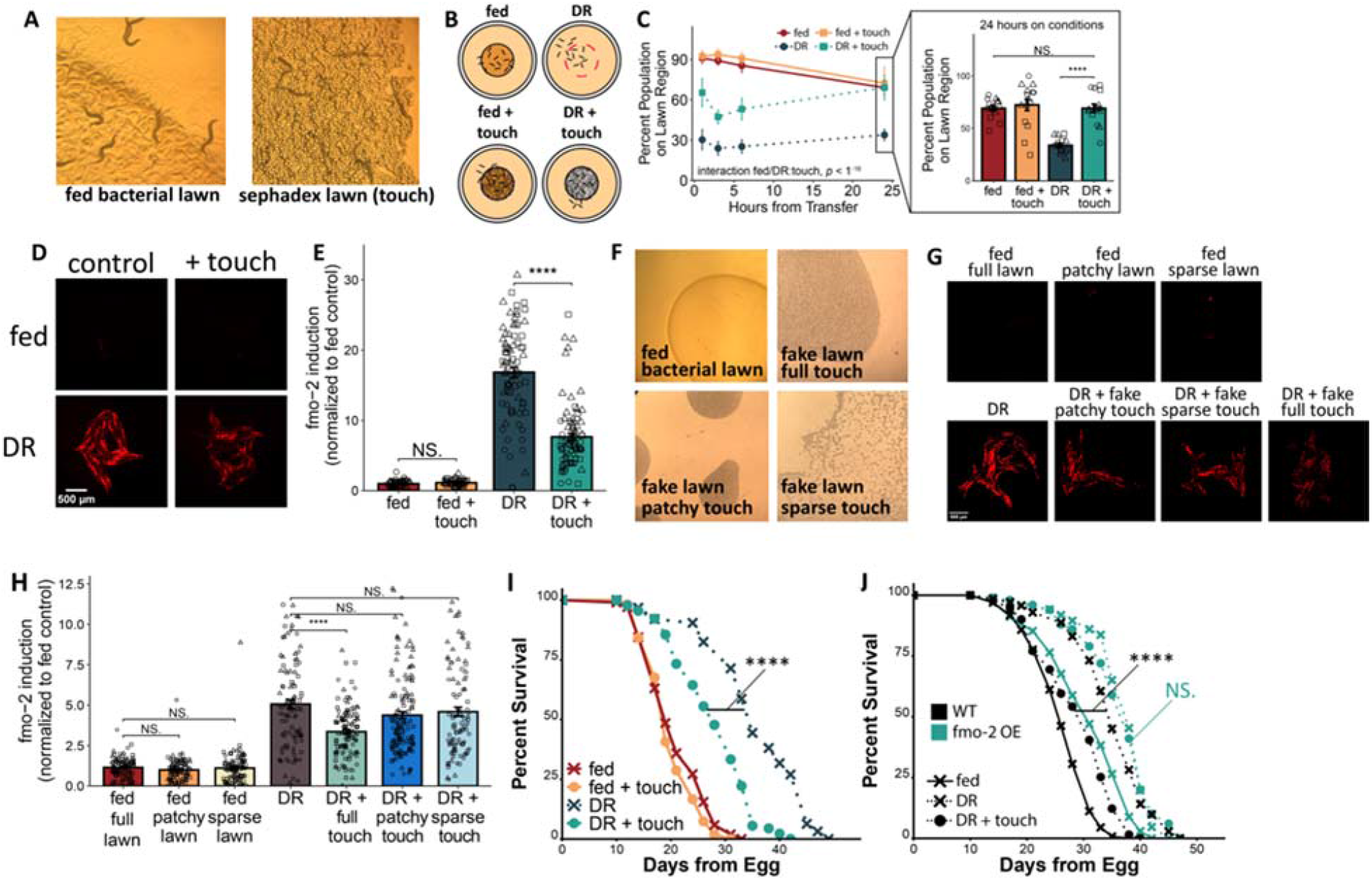
Food touch blunts DR-mediated longevity through suppression of *fmo-2*. (**A**) Images of *C. elegans* in real and fake bacterial lawns. (**B**) Schematic of fed/DR +/- touch conditions. (**C**) Lawn occupancy on fed/DR +/- touch conditions over time. *N* ≥ 45 worms/condition. (**D-E**) Representative images (**D**) and quantification (**E**) of *fmo-2p::mCherry* on fed/DR +/- touch. *N* ≥ 49 worms/condition. (**F**) Images of full, patchy, and sparse lawn configurations. (**G-H**) Representative images (**G**) and quantification (**H**) of *fmo-2p::mCherry* on fed/DR +/- different lawn configurations. *N* ≥ 99 worms/condition. (**I-J**) Survival curves of WT (**I**) and fmo-2 OE (**J**) animals on fed/DR +/- touch. *N* ≥ 63 worms/condition. (**C**,**E**,**H**) Show mean +/- SEM. Shapes indicate 3 biological replicates. Wilcox rank sum test with Bonferroni p-adjustment, two-sided, unpaired. (**I-J**) log rank test. In all panels, NS. = *p* > 0.05, ^****^ = *p*□< □.0001.

Based on our previous finding that food smells can decrease DR-mediated longevity by modulating the longevity gene *fmo-2* ^8^, we next asked whether food-like touch would blunt DR-mediated *fmo-2* induction. By imaging a single-copy integrated transcriptional *fmo-2* reporter under fed and DR conditions with and without a fake lawn (**Fig. 1D-E**), we found that touch did indeed decrease *fmo-2* induction under DR without affecting fed worms. Additionally, we confirmed that this *fmo-2* blunting effect is not due to a chemosensory property of the beads, as worms exposed to the supernatant of beads in water did not exhibit decreased *fmo-2* induction (**Fig. S1D-E**).

Next, we asked whether the beads composing the fake lawn had to be in a dense and lawn-like formation to suppress DR-mediated *fmo-2* induction. To answer this question, we measured *fmo-2* induction on DR alone, DR with the fake lawn, DR with a sparse lawn (equal mass of beads distributed in a thin layer), and DR with a patchy lawn (equal mass and concentration of beads distributed in small patches) (**Fig. 1F-H**). We found that only the full fake lawn decreased *fmo-2* induction under DR, establishing that the density and distribution of the fake lawn is crucial in decreasing the DR response. Finding that a dense fake lawn blunts expression of the longevity effector *fmo-2* suggests that touch may also blunt the healthspan and lifespan benefits of DR. After conducting a lifespan assay on the same four conditions, we found that the fake lawn does blunt DR-mediated longevity (**Fig. 1I**) but does not shorten fed lifespan. Moreover, this DR-blunting effect was not observed in worms that genetically overexpress FMO-2 (**Fig. 1J**), suggesting that decreased *fmo-2* induction due to mechanosensation causally suppresses DR-mediated longevity.

Having identified that the fake lawn blunts DR-mediated longevity in an *fmo-2*-dependent manner, we asked whether canonical mechanoreceptors are required. *C. elegans* respond to touch through >40 primary mechanoreceptors distributed throughout the body wall and nose tip ^13^. Both classes of *C. elegans* mechanoreceptors—transient receptor potential (TRP) and degerin/epithelial (DEG/ENaC) receptors—are also encoded in mammals ^14, 15^. To test whether broad inhibition of mechanosensation prevents the fake lawn from blunting DR, we put *fmo-2* reporters on fed, DR, and DR + touch conditions with and without amiloride, a DEG/ENaC inhibitor ^16, 17^. We found that 250μM of amiloride consistently blocked our phenotype (**Fig. 2A-B**), and that lower doses (10μM and 100μM) sometimes blocked the effect, with lower consistency across replicates (**Fig. S2A-B**). 250μM amiloride also prevented touch from abrogating DR-mediated longevity (**Fig. 2C**), while 100μM partially blocked the phenotype and 10μM had no effect on lifespan (**Fig. S2C-D**).

**Fig. 2.**
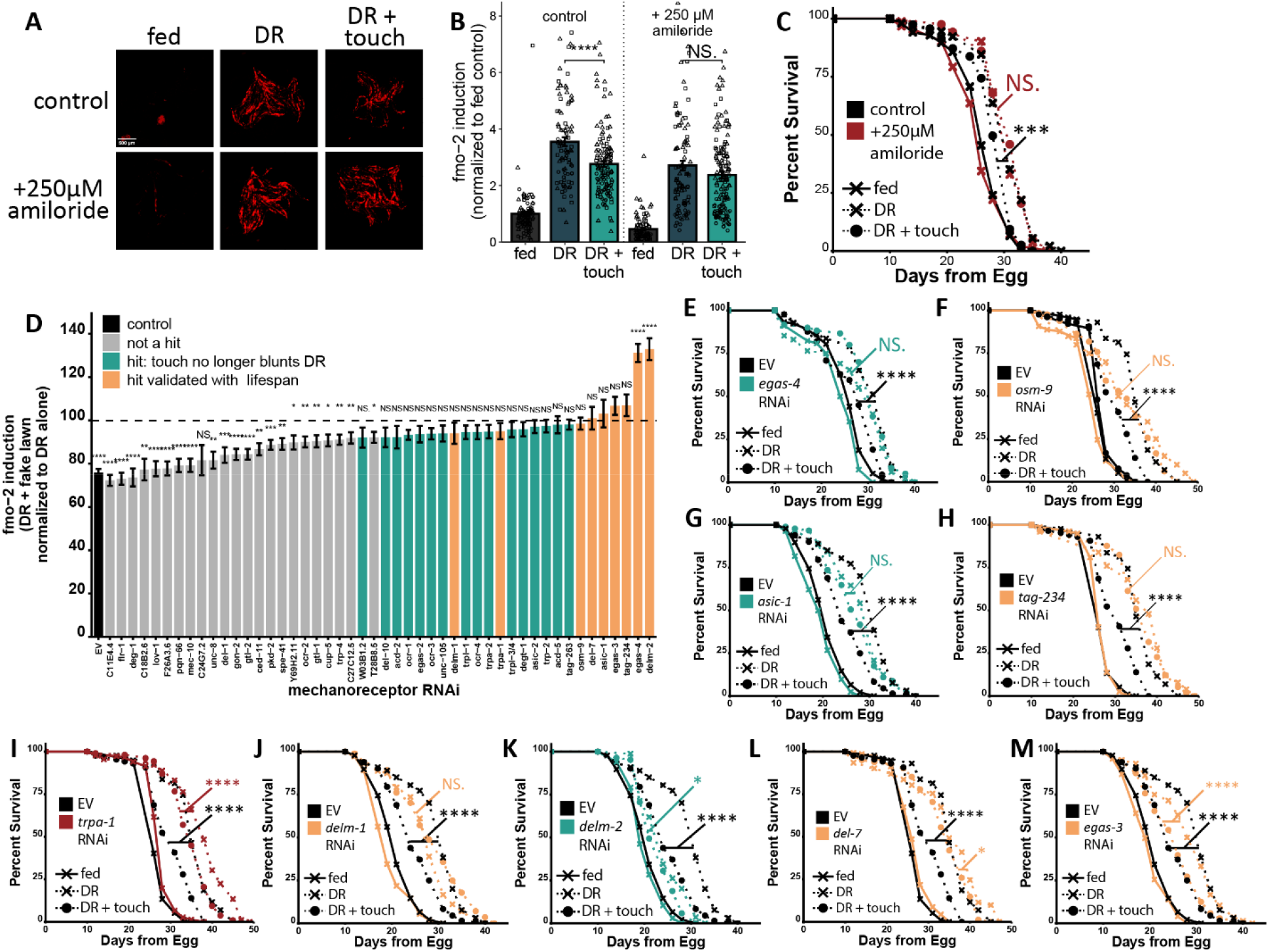
Multiple primary mechanoreceptors affect the ability of the fake lawn to blunt DR-mediated longevity. (**A-B**) Representative images (**A**) and quantification (**B**) of *fmo-2p::mCherry* on fed/DR +/- touch, +/- 250μM amiloride. *N* ≥ 73 worms/condition. Shapes indicate 3 biological replicates. (**C**) Survival curve of WT animals on fed/DR +/- touch conditions +/- 250μM amiloride. *N* ≥ 82 worms/condition. (**D**) Quantification of *fmo-2* induction on DR + touch as a percent of DR for each mechanoreceptor knockdown. *N* ≥ 68 for DR alone and ≥ 64 for DR + touch. (**E-M**) Survival curves of MAH677 on EV or (**E**) *egas-4* (*N* ≥ 43), (**F**) *osm-9* (*N* ≥ 20), (**G**) *asic-1* (*N* ≥ 82), (**H**) *tag-234* (*N* ≥ 25), (**I**) *trpa-1* (*N* ≥ 59), (**J**) *delm-1* (*N* ≥ 132), (**K**) *delm-2* (*N* ≥ 43), (**L**) *del-7* (*N* ≥ 72), or (**M**) *egas-3* (*N* ≥ 43) RNAi, on fed/DR +/- touch conditions. (**B, D**) Bar height shows mean +/- SEM. Wilcox rank sum test with Bonferroni p-adjustment, two-sided, unpaired. (**E-M**) Log rank test. In all panels, NS. = *p* > 0.05, ^*^ = *p* < 0.05, ^**^ = *p* < 0.01, *** = *p* < 0.001, ^****^ = *p*□< □0.0001.

After finding that DEG/ENaC receptors are required for touch to attenuate DR-mediated longevity, we wondered whether TRP channels also contribute, and whether any single mechanoreceptor is necessary. To answer these questions, we used RNAi to knock down every putative *C. elegans* mechanoreceptor with an available RNAi clone and measured *fmo-2* induction on fed and DR +/- touch. We found that knockdown of 25 mechanoreceptors prevented touch from blunting DR-mediated *fmo-2* induction (**Fig. 2D**) and many knockdowns altered baseline *fmo-2* induction without touch (**Fig. S3A-B**). Interestingly, many of these hits also modulate sensory modalities other than mechanosensation. OSM-9, for example, is also important in chemosensation and temperature sensation ^18^ while TRPA-1 plays a role in nociception and temperature perception ^19-21^.

Based on our imaging screen results, we followed up on nine mechanoreceptor hits that 1) blocked the DR + touch effect most, 2) were consistent across all biological replicates, and 3) included hits from both the DEG/ENaC and the TRP receptor categories. After conducting lifespan assays, we found that all nine knockdowns either completely (**Fig. 2E-J**) or partially (**Fig. 2K-M**) prevented touch from blunting DR (log rank test DR vs DR + touch and cox regression for an interaction between RNAi and touch, **Fig. S3C**).

### Signaling from dopamine and tyramine is required for mechanosensation to regulate longevity

Upon identifying multiple primary mechanoreceptors that regulate touch attenuation of DR, we sought to determine the neural signals downstream of initial mechanosensory perception. Identifying these signals could point to neural regulators of longevity downstream of sensory perception. To interrogate these signals, we crossed our *fmo-2* reporter with mutants deficient in the synthesis or packaging of each *C. elegans* neurotransmitter ^22^. We found that knocking out dopamine synthesis prevented touch attenuation of DR-mediated *fmo-2* induction (**Fig. 3A-C**).

**Fig. 3.**
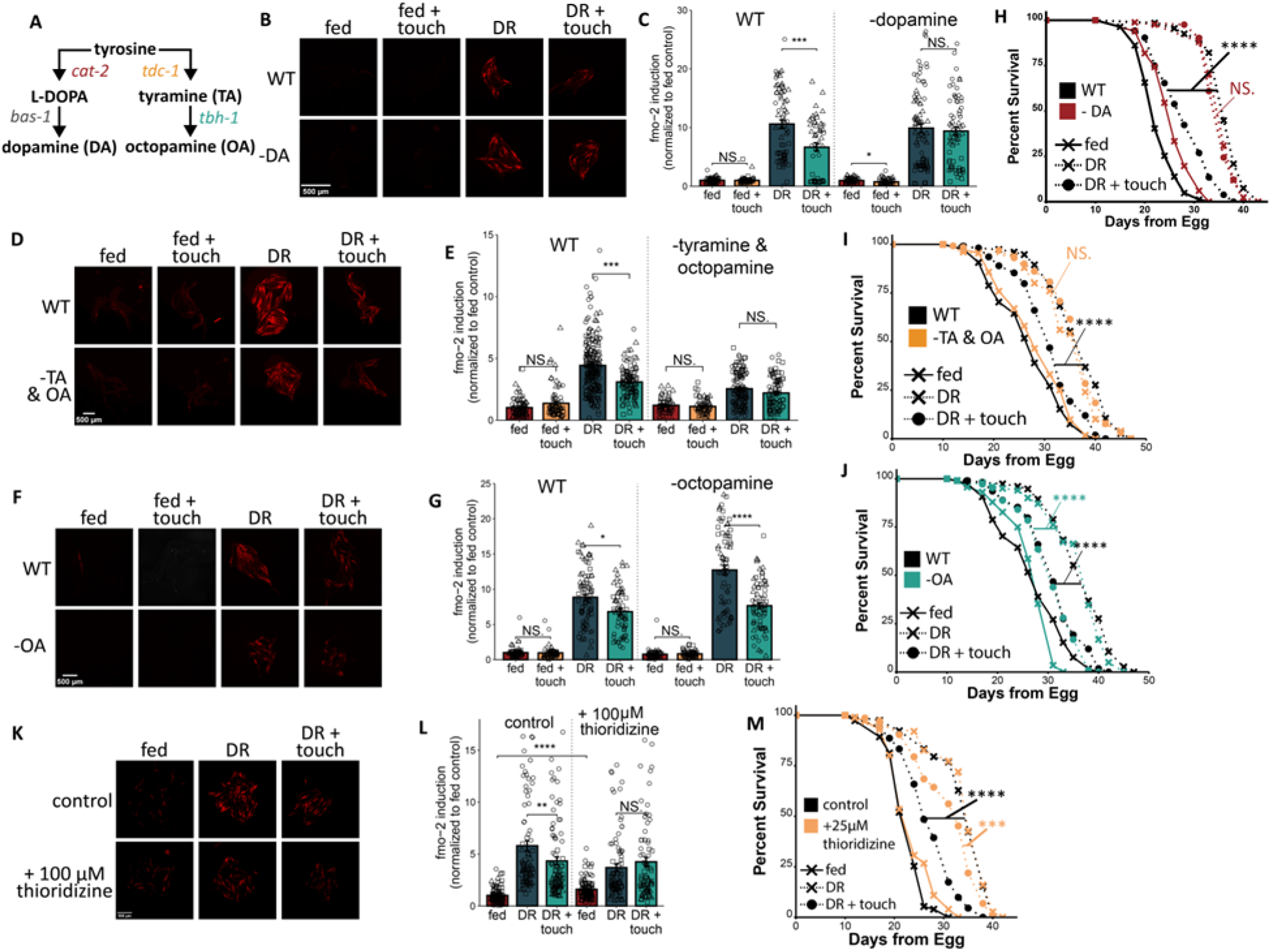
The bioamines dopamine and tyramine modulate DR-mediated longevity downstream of food touch. (**A**) Diagram of dopamine, tyramine, and octopamine synthesis. (**B-G**) Representative images (**B**,**D**,**F**) and quantification (**C, E, G**) of *fmo-2p::mCherry* (WT) and *fmo-2p::mCherry; cat-2(n4547)* (-dopamine, **B-C**), *fmo-2::mCherry; tdc-1(n3419)* (-tyramine & octopamine, **D-E**), or *fmo-2p::mCherry; tbh-1(n3247)* (-octopamine, **F-G**) on fed/DR +/- touch conditions. *N* ≥ 50 (**C**), ≥ 54 (**E**), and ≥ 54 (**G**) worms/condition. (**H-J**) Survival curves of WT and *cat-2(n4547)* (-DA) (**H**), *tdc-1(n3419)* (-TA & OA) (**I**), or *tbh-1(n3247)* (-OA) (**J**) animals on fed/DR +/- touch conditions. *N* ≥ 41 (**H**), ≥ 50 (**I**), ≥ 76 (**J**) per condition. (**K-L**) Images (**K**) and quantification (**L**) of *fmo-2p::mCherry* on fed/DR +/- touch with or without 100μM thioridazine. *N* ≥ 77 worms per condition. (**M**) Survival curves of WT animals on fed/DR +/- touch conditions with and without 25μM thioridazine. *N* ≥ 73 worms per condition. (**C, E, G, L**) Bar height shows mean +/- SEM. Wilcox rank sum test with Bonferroni p-adjustment, two-sided, unpaired. Shapes indicate biological replicates. (**H-J, M**) Log-rank test. In all panels, NS. = *p* > 0.05, ^*^ = *p* < 0.05, ^**^ = *p* < 0.01, *** = *p* < 0.001, ^****^ = *p*□< □0.0001.

Additionally, knocking out tyramine and octopamine synthesis blocked the touch phenotype (**Fig. 3A, D-E**). However, knocking out octopamine synthesis alone (**Fig. 3A, F-G**) had no effect on the touch response, suggesting that tyramine but not octopamine is required for this effect.

Impairing the synthesis or packaging of serotonin (**Fig. S4A-B**), GABA (**Fig. S4C-D**), glutamate (**Fig. 4SE-F**), and acetylcholine (**Fig. S4G-H**) also had no effect on the ability of touch to blunt DR. The same effects were observed in lifespan assays: blocking dopamine synthesis (**Fig. 3H**) and blocking tyramine and octopamine synthesis (**Fig. 3I**) also prevented touch from attenuating DR-mediated longevity while blocking octopamine synthesis alone (**Fig. 3J**) did not.

**Fig. 4.**
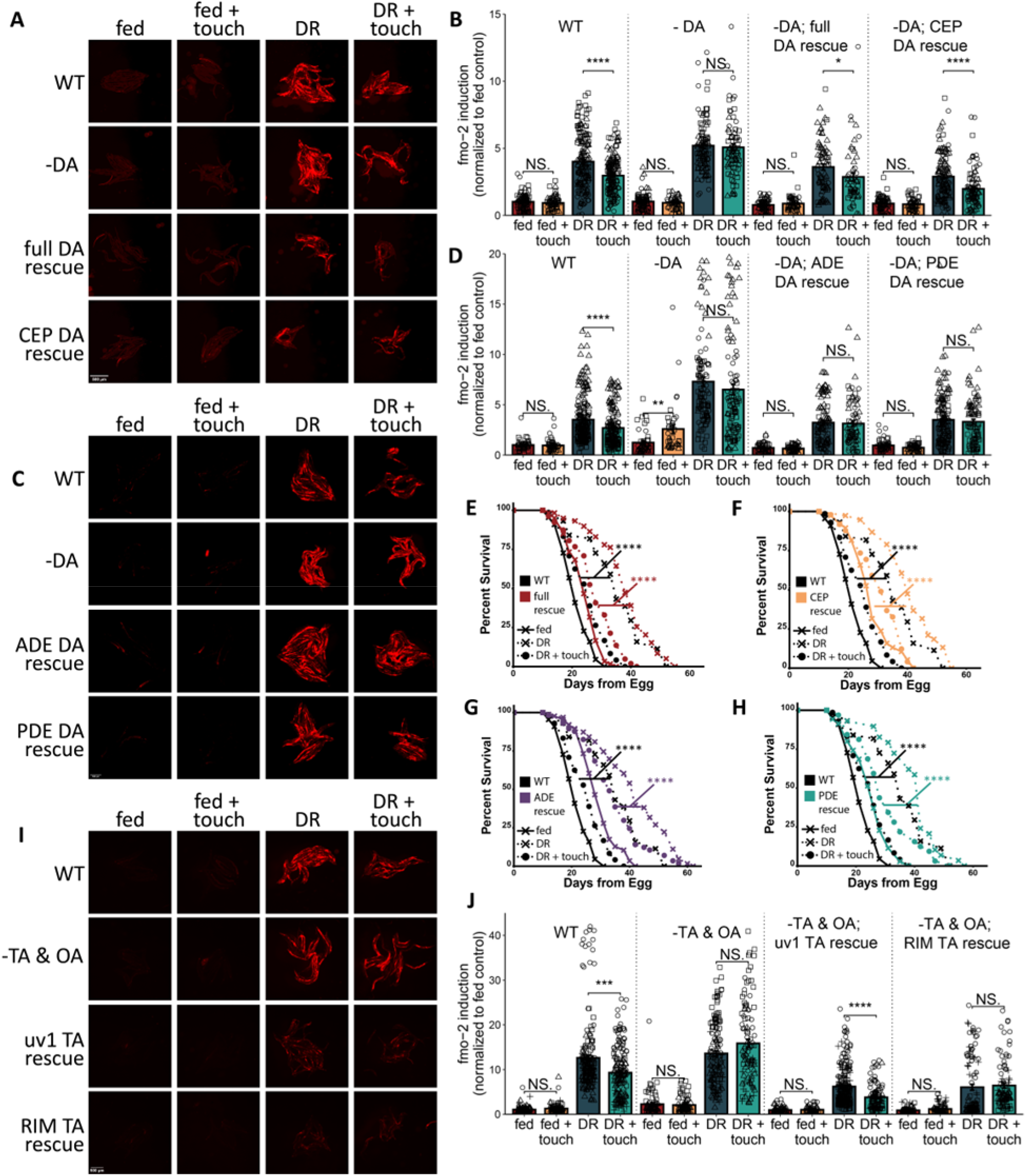
Dopamine production from any dopaminergic neuron class and tyramine production from the uv1 neuroendocrine cells is sufficient for food touch to blunt DR. (**A-B**) Representative images (**A**) and quantification (**B**) of *fmo-2p::mCherry* (WT), *fmo-2p::mCherry; cat-2(n4547)* (-DA), *fmo-2p::mCherry; cat-2(n4547); Ex[cat-1p::cat-2]* (full DA rescue) and *fmo-2p::mCherry; cat-2(n4547);Ex[swip-10p::cat-2]* (CEP DA rescue) on fed/DR +/- touch conditions. *N* ≥ 51 worms per condition. (**C-D**) Representative images (**C**) and quantification (**D**) of *fmo-2p::mCherry* (WT), *fmo-2p::mCherry cat-2(n4547)* (-DA), *fmo-2p::mCherry; cat-2(n4547);Ex[trpa-1p::cat-2]* (ADE DA rescue) and *fmo-2p::mCherry; cat-2(n4547); Ex[tax-2p::cat-2]* (PDE DA rescue) on fed/DR +/- touch conditions. *N* ≥ 37 worms per condition. (**E-H**) Survival curves of WT and *cat-2(n4547); Ex[cat-1p::cat-2]* (full DA rescue, **E**), *cat-2(n4547);Ex[swip-10p::cat-2]* (CEP DA rescue, **F**), *cat-2(n4547);Ex[trpa-1p::cat-2]* (ADE DA rescue, **G**), or *cat-2(n4547); Ex[tax-2p::cat-2]* (PDE DA rescue, **H**) animals on fed/DR +/- touch conditions. *N* ≥ 44 (**E**), ≥ 38 (**F**), ≥ 44 (**G**), and ≥ 26 (**H**) per condition. (**I-J**) Representative images (**I**) and quantification (**J**) of *fmo-2p::mCherry* (WT), *fmo-2p::mCherry; tdc-1(n3419)* (-TA & OA), *fmo-2p::mCherry; tdc-1(n3419); Ex[ocr-1p::tdc-1]* (uv1 TA rescue), and *fmo-2p::mCherry; tdc-1(n3419); Ex[gcy-13p::tdc-1]* (RIML TA rescue) on fed/DR +/- touch conditions. *N* ≥ 66 worms per condition. (**B, D, J**) Bar height shows mean +/- SEM. Wilcox rank sum test with Bonferroni p-adjustment, two-sided, unpaired. Shapes indicate biological replicates. (**E-H**) Log-rank test. In all panels, NS. = *p* > 0.05, ^*^ = *p* < 0.05, ^**^ = *p* < 0.01, *** = *p* < 0.001, ^****^ = *p*□< □ 0.0001.

Interestingly, the production of both dopamine and tyramine (the invertebrate analog of adrenaline) is regulated by food availability. Dopamine release is generally associated with finding food ^12, 23^, but the role of tyramine in signaling nutrient status is less clear ^24, 25^. When we supplemented *fmo-2* reporters with exogenous dopamine, we observed a dose-dependent decrease in DR-mediated *fmo-2* induction (**Fig. S4I-J**). Exogenous tyramine, on the other hand, dose-dependently increased *fmo-2* induction on fed conditions (**Fig. S4K-L**). This suggests a model in which touch perception triggers dopamine release and suppresses tyramine release to attenuate DR-mediated longevity.

In our previous work on the role of olfaction in DR, we found that two small molecules that antagonize dopamine and serotonin (thioridazine and mianserin, respectively), act as DR-mimetics to extend lifespan ^8^. Consistent with our neurotransmitter screen results, we found that thioridazine (**Fig. 3K-M**) but not mianserin (**Fig. S4M-O**) prevented the fake lawn from blunting DR-mediated *fmo-2* induction (**Fig. 3K-L, S4M-N**) and longevity (**Fig. 3M, S4O**). These results indicate that DR mimetics acting downstream of food perception may function regardless of external nutrient cues. Moreover, the ability of thioridazine but not mianserin to block food touch further supports a role for dopamine, but not serotonin, as an intermediate signal in multiple circuits through which perception regulates DR-mediated longevity.

Once we established the necessity of dopamine and tyramine, we asked which dopaminergic and tyraminergic cells were key players in the DR touch circuit. To determine which dopaminergic neuron(s) signal in this pathway, we rescued dopamine production in each of the three *C. elegans* dopaminergic neuron classes in a dopamine deficient background strain. We found that rescuing dopamine production in all dopaminergic cells and just in the CEP neurons restored WT-like *fmo-2* induction (**Fig. 4A-B**) while rescuing dopamine in the ADE or PDE neurons did not (**Fig. 4C-D**). Unlike in our imaging experiments, however, we found that rescuing dopamine production in any dopaminergic cell type rescued the effect of touch on DR-mediated longevity (**Fig. 4E-H**). Together, these results are consistent with a model in which a subpopulation of dopaminergic neurons play a key role in touch modification of *fmo-2* induction, while neuronal specificity does not contribute to touch modification of lifespan. It is also notable that signaling from all three classes of dopaminergic neurons are required for worms to slow upon encountering a bacterial lawn ^12^, suggesting a complex interaction between the localization of dopamine signaling and various physiological responses to food cues.

In addition to investigating key dopaminergic neurons, we also asked whether signaling from a specific tyraminergic neuron type was sufficient to restore WT responses to DR + touch. In *C. elegans*, the RIML neuron and uv1 neurodendocrine cells are considered tyraminergic because they express *tdc-1* but not *tbh-1*^*26, 27*^ (**Fig. 3A**). When we rescued *tdc-1* expression to restore tyramine synthesis in each of these cell types, we found that only the uv1 cell rescue was sufficient to restore WT-like *fmo-2* induction in response to DR and touch (**Fig. 4I-J**). These data suggest a model in which mechanosensation modulates tyramine release from the uv1 neuroendocrine cells and dopamine release from all dopaminergic neurons, which ultimately send a signal to the intestine to blunt DR-mediated *fmo-2* induction and longevity.

### Insulin-like neuropeptide signaling conveys touch information from the nervous system to the intestine

While many dopamine- and tyramine-producing neurons are immediately downstream of primary mechanosensory neurons ^28^, our data suggest it is unlikely that bioaminergic cells signal to the intestine to suppress *fmo-2*. The *C. elegans* nervous system does not directly innervate peripheral tissues and worms instead secrete neuropeptide/neuroendocrine signals into the pseudocoelom to communicate with these tissues ^29^. To examine whether neuropeptide signaling may facilitate this cell-nonautonomous touch signal, we examined the DR + touch response of the dense-core vesicle packing mutant *unc-31* ^*30*^. We found that the fake lawn did not suppress DR-mediated *fmo-2* induction or longevity in *unc-31* mutants (**Fig. 5A-C**). We also found that knocking down the proneuropeptide maturation and cleavage genes *egl-3* and *egl-21* ^29^ prevented touch from attenuating DR-mediated *fmo-2* induction (**Fig. S5A-B**). Together, these results suggest a role for neuropeptides in conveying mechanosensory information from the nervous system to intestinal cells that express the DR effector protein, FMO-2.

**Fig. 5.**
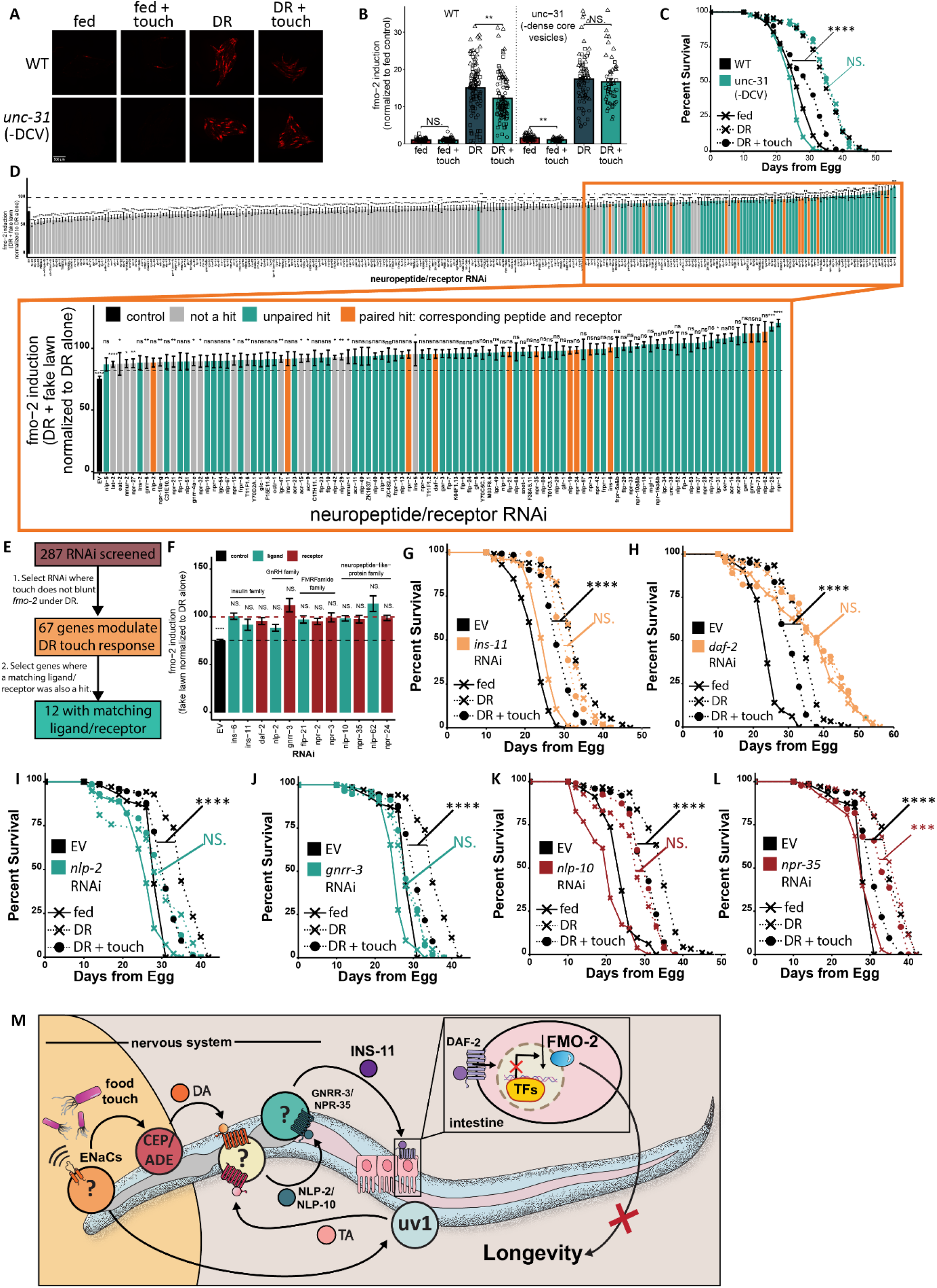
Insulin-, GnRH-, and NLP-family neuroendocrine signals are required for touch to blunt DR. (**A-B**) Representative images (**A**) and quantification (**B**) of *fmo-2p::mCherry* (WT) and *fmo-2p::mCherry; unc-31* (unc-31) on fed/DR +/- touch conditions. *N* ≥ 51 worms per condition. Shapes indicate 3 biological replicates. (**C**) Survival curves of WT and *unc-31* worms on fed/DR conditions +/- touch. *N* ≥ 49 worms per condition. (**D**) Quantification of *fmo-2p::mCherry; TU3311* on DR + touch as a percent of DR for each neurosignaling knockdown. *N* ≥ 54 for DR alone and ≥ 56 for DR + touch. (**E**) Flow diagram for selecting top hits. (**F**) *fmo-2* induction on DR + touch as a percent of DR alone on each of the 12 paired hits from (**D**). (**G-L**) Survival curves of TU3311 on EV or *ins-11* (**G**), *daf-2* (**H**), *nlp-2* (**I**), *gnrr-3* (**J**), *nlp-10* (**K**), and *npr-35* (**L**) RNAi, on fed/DR +/- touch conditions. *N* ≥ 73 (**G**), ≥ 44 (**H**), ≥ 37 (**I**), ≥ 51 (**J**), ≥ 44 (**K**), ≥ 44 (**L**) per condition. (**M**) Diagram of mechanosensory modulation of aging under DR conditions. (**B, D, F**) Bar height shows mean +/- SEM. Wilcox rank sum test with Bonferroni p-adjustment, two-sided, unpaired. (**G-L**) Log rank test. In all panels, NS. = *p* > 0.05, ^*^ = *p* < 0.05, ^**^ = *p* < 0.01, *** = *p* < 0.001, ^****^ = *p*□< □0.0001.

To identify neuropeptide candidates for this cell-nonautonomous signal, we used RNAi to knock down 287 potential neurosignaling ligands and receptors in our *fmo-2* reporter.

Knockdowns resulting in no significant difference between *fmo-2* induction on DR and DR + touch were considered hits. Based on these initial criteria, we identified 85 neurosignaling genes that modified the DR-touch response (**Fig. 5D**). To narrow down this set of potential hits, we eliminated RNAi where results were not consistent across at least 2/3 of replicates. This left us with 67 consistent hits, which we further prioritized by identifying ligand/receptor pairs in which both the ligand and a receptor known to bind to that ligand blocked the DR-touch effect. This final criterion left us with a set of 12 genes falling into 5 groups of ligand/receptor matches (**Fig. 5E-F**).

When we followed up on these 12 hits with lifespans (all touch effects summarized in **Fig. S5C**), we found that 8 of the 12 RNAi imaging hits completely (**Fig. 5G-K, Fig. S5D-E**) or partially (**Fig. 5L**) prevented touch from attenuating DR, while the remaining four did not validate (**Fig. S5F-I**). Of the five neuropeptide ligand/receptor families tested, both members of a matching ligand/receptor pair blocked the DR + touch effect in the insulin family (*daf-2, ins-11*), the GnRH family (*nlp-2, gnrr-3*), and one of the two neuropeptide-like protein families (*nlp-10* full block, *npr-35* partial block). Notably, DAF-2 is the only one of these receptors known to be expressed in the intestine (**Fig. S5J**) ^31^, suggesting that *ins-11/daf-2* signaling could be the cell-nonautonomous signal conveying mechanosensory information to the intestine. To further validate the role of insulin signaling in the touch-DR pathway, we crossed our *fmo-2* reporter with a mutant of *daf-16*, a key transcription factor acting downstream of DAF-2. We found that, like *ins-11* and *daf-2, daf-16* was required for touch to blunt *fmo-2* induction under DR (**Fig S5K-L**). Interestingly, previous work identified a role for *ins-11* in cell-nonautonomous signaling from the intestine to the nervous system in response to pathogenic bacteria ^32^.

While INS-11/DAF-2 signaling is a likely candidate for communication from the nervous system to the intestine, *nlp-2*/GnRH and *nlp-10* signals may integrate touch information within the nervous system and/or germline. Interestingly, *nlp-2*/GnRH is highly expressed in the tyraminergic uv1 cells ^33^, suggesting a potential integration point between the bioamine and neuropeptide branches of this circuit. Hypothalamic GnRH release is also a direct regulator of reproductive hormones in mammals, and consequently is highly sensitive to nutrient availability^34^. A role for GnRH signaling in aging would also be consistent with the reproductive/longevity tradeoff in which organisms can either conserve resources for maximum longevity or devote resources to maximize reproduction ^35^. Taken together, our data support a model in which gentle touch activates multiple mechanoreceptors that modulate signaling from dopaminergic neurons and tyraminergic uv1 cells. These bioamines then modify neuroendocrine signaling through NLP-2/GnRH, NLP-10, and INS-11. Ultimately, this food-touch circuit decreases intestinal *fmo-2* induction and longevity under DR conditions (**Fig. 5M**). These findings support a growing literature on the role of sensory perception in healthy aging. Under dietary restriction conditions specifically, food-like smells ^6-8^ and tastes ^9, 10^ attenuate DR-mediated longevity. In this work, we identify a role for mechanosensation in DR and connect sensory perception to known (bioamines, insulin, FMO-2) and novel (GnRH, NLP-10) longevity-modifying signals.

While technically challenging, future work in this area should focus on identifying interneurons acting downstream of multiple modes of sensory perception that modulate aging. Currently, it remains unclear whether discrete environmental stimuli that regulate aging act through convergent or parallel pathways to modify lifespan. Additionally, it is notable that perception of food-like smell, taste, or touch alone is not sufficient to completely block the DR response. This suggests that either 1) different modes of food perception are integrated under fed conditions to completely suppress a DR response, or 2) metabolic changes that occur under nutrient scarcity are also accounted for when determining the magnitude of the DR-response.

Examining the interplay between metabolic and sensory signals of nutrient scarcity could lead to key insights regarding the maximum effect sensory perception can have on longevity.

