## Supplementary material for "Food touch limits lifespan through bioamine and neuroendocrine signaling": Methods and Supplemental Information

**Supplementary Information/Extended Data**

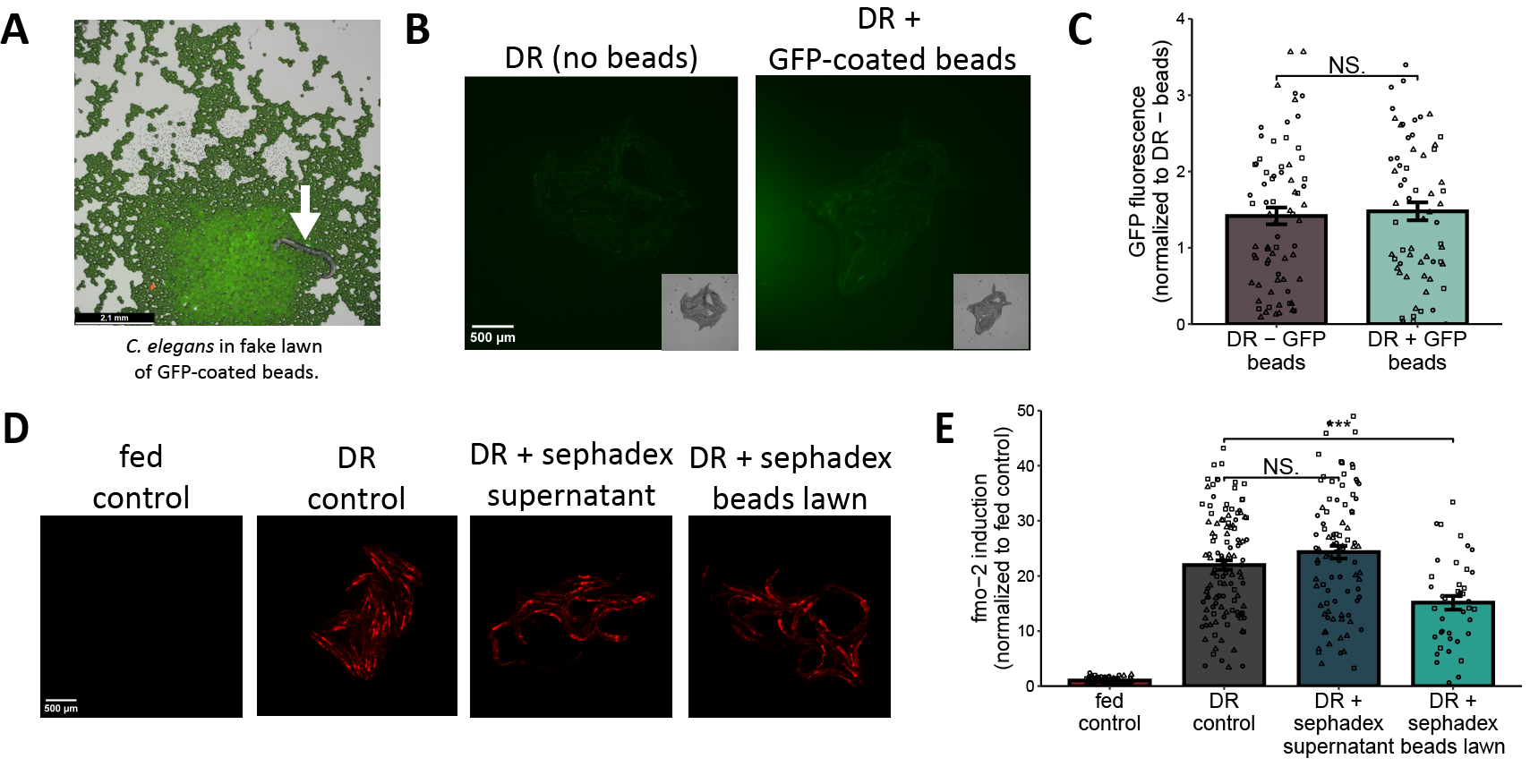

**Fig. S1. The fake lawn is not edible and does not blunt DR-mediated *fmo-2* induction via chemosensation.**

(**A**) *C. elegans* (white arrow) among GFP-coated 20-50μM beads. (**B-C**) Images (**B**) and quantification (**C**) of WT N2 worms on a DR +/- GFP-coated beads. *N* ≥ 64 worms. **D-E)** Images **(D)** and quantification **(E)** of *fmo-2p::mCherry* on fed and DR control conditions as well as DR + 200μL sephadex media supernatant and DR + 200μL sephadex beads for 24 hours. *N* ≥ 71 worms. Quantification in (**C, E**) shows mean +/- SEM. Shapes indicate biological replicates. Wilcox rank sum test with Bonferroni p-adjustment, two-sided, unpaired, NS. = *p* > 0.05, *** = *p* < .001.

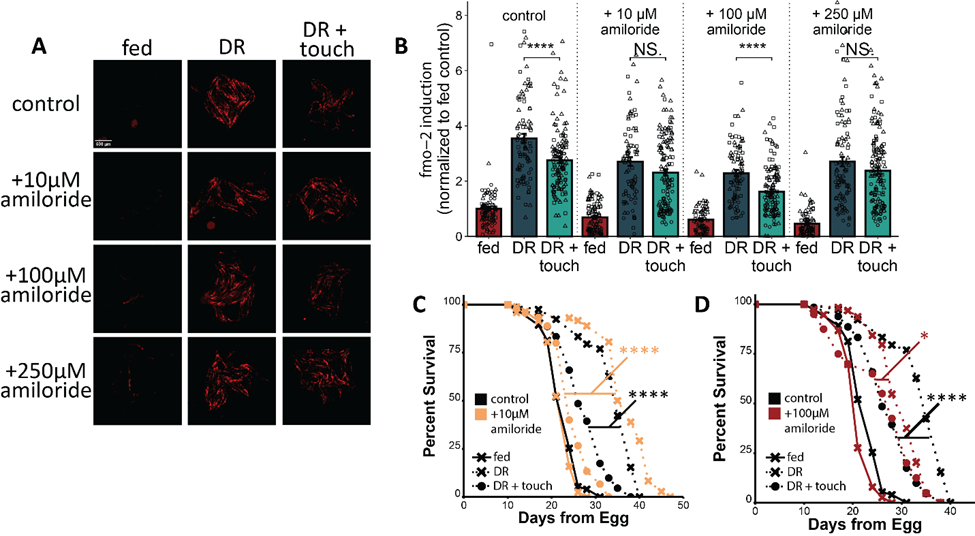
 Fig. S2. 250μM amiloride has the greatest blocking effect on DR + touch.

(**A-B**) Representative images (**A**) and quantification (**B**) of *fmo-2p::mCherry* on fed/DR +/- touch, +/- 10 μM, 100 μM, or 250μM amiloride. *N* ≥ 73 worms/condition. Bars show mean +/- SEM. Shapes indicate biological replicates. Wilcox rank sum test with Bonferroni p-adjustment, two-sided, unpaired. (**C-D**) Survival curve of WT animals on fed/DR +/- touch conditions +/- 10μM (**C**) or 100μM (**D**) amiloride. *N* ≥ 45 (**C**) or *n* ≥ 40 (**D**) worms/condition. Log-rank tests. In all panels, NS. = *p* > 0.05, * = *p* < 0.05, **** = *p* < 0.0001.

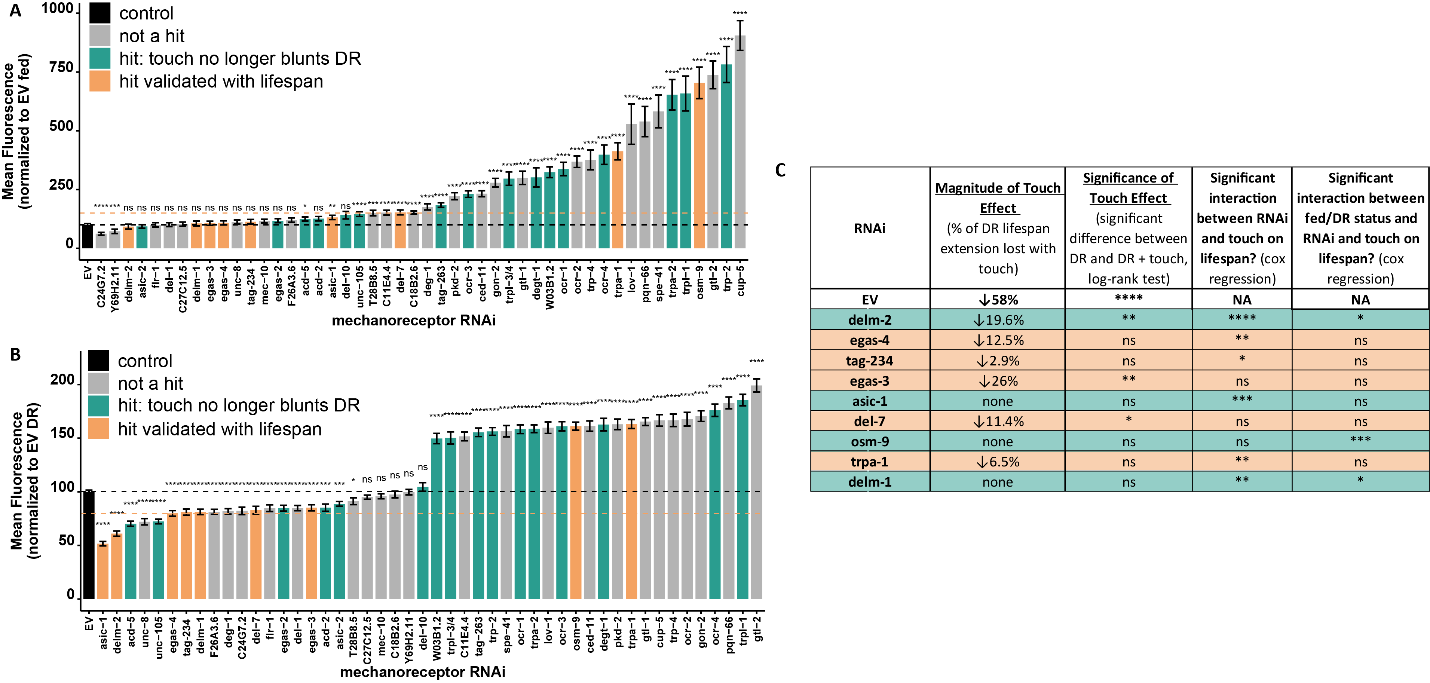
**Fig. S3. Knocking down many mechanoreceptors alters baseline fed and DR *fmo-2* induction and longevity with and without touch.**

(**A-B**) Quantification of *fmo-2p::mCherry* on fed (**A**) or DR (**B**) as a percent of the EV control for each mechanoreceptor knockdown. *N* ≥ 47 (**A**) or *n* ≥ 68 (**B**). Bar height shows mean of 3 biological replicates +/- SEM. Wilcox rank sum test with Bonferroni p-adjustment, two-sided, unpaired. (**C**) Effect of top imaging hits on DR + touch lifespan. In all panels, NS. = *p* > 0.05, * = *p* < 0.05, ** = *p* < 0.01, *** = *p* < 0.001, **** = *p* < 0.0001.

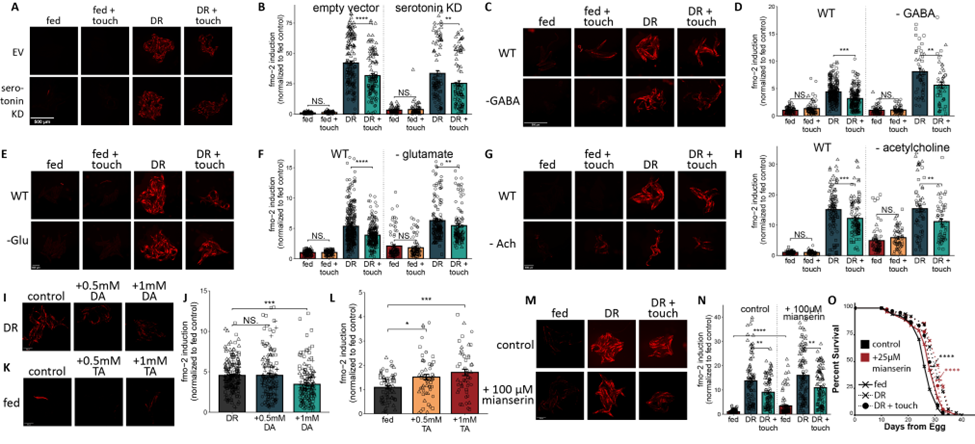

Fig. S4. Dopamine and tyramine signaling, but not serotonin, GABA, glutamate, or acetylcholine signaling are required for touch to attenuate DR.

(**A-B**) Representative images (**A**) and quantification (**B**) of *sid-1(qt9); sqIs71; fmo-2p::mCherry* on empty vector or *tph-1* RNAi (serotonin KD) on fed/DR +/- touch conditions. *N* ≥ 61 worms per condition. (**C-H**) Representative images (**C, E, G**) and quantification (**D, F, H**) of *fmo-2p::mCherry* (WT) and *fmo-2p::mCherry; unc-25(e156)* (-GABA, **C-D**), *fmo-2::mCherry; eat-4(ky5)* (-glutamate, **E-F**), or *fmo-2p::mCherry; unc-17(e113)* (-acetylcholine, **G-H**) on fed/DR +/- touch conditions. *N* ≥ 50 (**D**), ≥ 75 (**F**), and ≥ 53 (**H**) worms/condition. (**I-L**) Representative images (**I, K**) and quantification (**J, L**) of *fmo-2p::mCherry* on DR (**I-J**) or fed (**K-L**) conditions with or without 0.5mM and 1mM exogenous dopamine (**I-J**) and tyramine (**K-L**). *N* ≥ 153 (**J**), and ≥ 52 (**L**) worms/condition. (**M-N**) Representative images (**M**) and quantification (**N**) of *fmo-2p::mCherry* on fed/DR +/- touch conditions with or without 100μM mianserin. *N* ≥ 91 worms per condition. (**O**) Survival curves of WT animals on fed/DR +/- touch conditions with and without 25μM mianserin. *N* ≥ 56 worms per condition. (**B, D, F, H, J, L, N**) Bar height shows mean +/- SEM. Wilcox rank sum test with Bonferroni p-adjustment, two-sided, unpaired. Shapes indicate biological replicates. (**O**) Log-rank test. In all panels, NS. = *p* > 0.05, * = *p* < 0.05, ** = *p* < 0.01, *** = *p* < 0.001, **** = *p* < 0.0001.

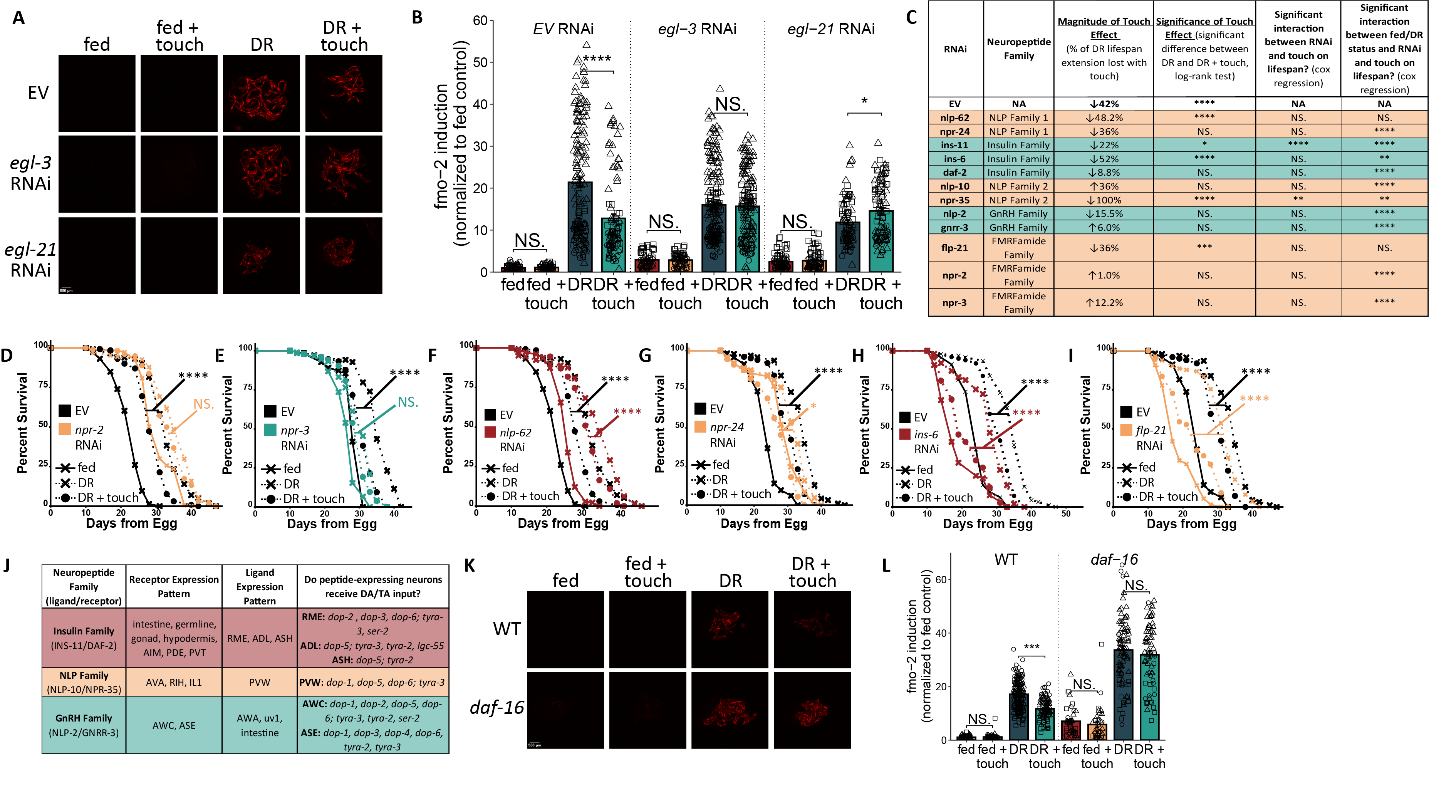
**Fig. S5. Many neuroendocrine genes regulate DR-mediated *fmo-2* induction and longevity within and independently of the touch pathway.**

Representative images (**A**) and quantification (**B**) of *fmo-2p::mCherry; MAH677* on fed/DR +/- touch conditions on EV, egl-3, or egl-21 RNAi. *N* ≥ 57 worms per condition. (**C**) Effect of top imaging hits on DR + touch lifespan. Log rank tests and cox regression tests with Bonferroni adjustment for comparison across 3 biological replicates. (**D-I**) Survival curves of TU3311 on EV or *npr-2* (**D**), *npr-3* (**E**), *nlp-62* (**F**), *npr-24* (**G**), *ins-6* (**H**), and *flp-21* (**I**) RNAi, on fed/DR +/- touch conditions. *N* ≥ 73 (**D**), ≥ 51 (**E**), ≥ 55 (**F**), ≥ 44 (**G**), ≥ 44 (**H**), ≥ 44 (**I**) per condition. (**J**) Summary of neurons and tissues that express each neuroendocrine signal hit, as well as whether these neurons receive input from dopamine and/or tyramine. (**K-L**) Representative images (**K**) and quantification (**L**) of *fmo-2p::mCherry* (WT), and *fmo-2p::mCherry; daf-16* (daf-16) on fed/DR +/- touch conditions. *N* ≥ 34 worms per condition. (**B-C, L**) Bar height shows mean +/- SEM. Wilcox rank sum test with Bonferroni p-adjustment, two-sided, unpaired. (**B, L**) Shapes indicate 3 biological replicates. (**D-I**) Log rank test. In all panels, NS. = *p* > 0.05, * = *p* < 0.05, ** = *p* < 0.01, *** = *p* < 0.001, **** = *p* < 0.0001.

**Materials and Methods**

Strains and Growth Conditions

*C. elegans* were maintained following standard procedures ^36^ in 20°C temperature-controlled incubators. Nematodes were fed *Escherichia coli* (OP50) seeded on solid nematode growth medium (NGM) plates. For maintenance and lifespans, worms were transferred gently with a platinum wire pick. A list of strains and RNAi conditions used in this study can be found in **Supplementary Tables 1 and 2,** respectively. All genotypes were confirmed using PCR and all RNAi imaging hits were sequence-validated before use in lifespans.

*tdc-1* Rescue Constructs

Plasmid construction and microinjection was conducted by Suny Biotech to generate both *tdc-1* rescue constructs. In designing the plasmids, we used the *ocr-4* promoter to drive cDNA of TDC-1::SL2::GFP in the uv1 neuroendocrine cells. To express cDNA of TDC-1::SL2::GFP in the RIML neuron, we used the *gcy-13* promoter. All plasmids were verified via restriction digest and sanger sequencing, and ApE files are available upon request. Plasmids were microinjected by Suny Biotech using the co-injection marker *myo-2*p::GFP and 3 transgenic lines were tested in each experiment.

Fake Lawn Preparation

G-200 Sephadex beads (Sigma G5050) were suspended S-basal buffer at a concentration of 30mg/ml and autoclaved for 45 minutes. For fed + touch plates, OP50 bacteria was grown overnight to an optical density (OD_600_) of 3.0, then pelleted at 3000 *rcf* for 20 minutes. The supernatant was discarded and replaced with the sephadex in S-basal, then the mixture was vortexed until the pellet was dissolved. 200µl was seeded onto the center of a 60mm diameter NGM plate. For DR + touch conditions, autoclaved sephadex in S-basal was added directly to the center of the NGM plate (overnight imaging experiments) or the autoclaved sephadex in S-basal was used to resuspend a pellet from a diluted concentration of bacteria (DR lifespan experiments). For fed controls and DR lifespan controls, the bacterial pellet was resuspended in S-basal instead of S-basal + sephadex. For DR imaging controls, 200µl autoclaved S-basal was added directly to the plate.

RNAi Knockdown

The RNAi feeding bacteria were grown from the Ahringer and Vidal *C. elegans* RNAi feeding libraries. All RNAi plasmids used for lifespan assays were sequenced to verify the correct target sequence. RNAi used for large screens (mechanoreceptor and neuroendocrine signaling screens) were not sequenced initially, but any hits that moved on to the validation stage were sequence-verified. Animals in RNAi knockdown conditions were progeny from two generations of adults on the RNAi. RNAi bacteria was seeded on NGM plates supplemented with 1 mM β-D-isothiogalactopyranoside (IPTG) and 25μg/ml carbenicillin. Previous work has found that the RNAi feeding bacteria (HT115 strain of *E. coli*) can alter worm lifespan relative to a diet of OP50 ^37, 38^. These differences should be noted when comparing lifespans conducted on RNAi relative to lifespans on OP50.

Drug Treatments

For all experiments with supplementation of exogenous small molecules (exogenous dopamine, exogenous tyramine, amiloride, thioridazine, and mianserin), the small molecule was dissolved in solid agar plates. Each compound was added to agar once it had cooled to < 55˚C and before it was poured into plates. All drugs were purchased from Sigma-Aldrich and were initially dissolved in milliQ water at 2mM (mianserin), or 100mM concentration (thioridazine, amiloride, dopamine, and tyramine), aliquoted, and stored at -20˚C.

Lifespan Measurements

Lifespan assays were conducted as previously described ^39^ with the exception of the fake lawn conditions (see fake lawn preparation). In short, 10-15 gravid adults were left on NGM plates for a 3-4 hour timed egg lay and then removed. Once the progeny of these adults reached late L4/early adult stage of development, they were transferred to NGM plates with 33µL of 150mM fluorodeoxyuridine (FUdR) and 100µL of 50mg/mL Ampicillin per 100mL NGM. The FUdR prevents development of progeny and the Ampicillin prevents bacterial growth. 60-80 worms were placed on each NGM + FUdR + AMP plate seeded with concentrated bacteria (5x for OP50 fed lifespans, 0.5x OP50 (10^9^ CFU/mL) for DR lifespans), and at least two plates per strain per condition were used for each lifespan replicate. In the DR and DR + touch conditions, worms were kept on 5x fed bacteria until day 2 of adulthood, when they were transferred to fresh DR or DR + touch plates every other day for four transfers. This form of DR is referred to as solid DR (sDR) ^40^. Worms on fed and fed + touch conditions were transferred to fresh fed plates at the same time as each DR transfer. Animals were removed from the experiment and counted as dead when they did not move in response to a gentle touch from a platinum wire pick under a dissection microscope. Lifespans were scored at least 3 times per week until all animals were dead. A ‘fence’ of 75µL of 100mM palmitic acid (Sigma-Aldrich) dissolved in 100% ethanol was also applied to the edge of each lifespan plate to prevent fleeing. While FUdR can extend lifespan, FUdR was used in all lifespan plates to prevent matricide under DR conditions ^41^. Data were plotted by R version 4.3.1 and Adobe Illustrator 2022.

RNAi Lifespans

RNAi lifespan assays were performed like other lifespans with the exception of the initial TEL and the food concentrations. To ensure maximum knockdown on RNAi, a first generation of worms was TEL’ed on RNAi for 3-4 hours. The progeny from this TEL were left on RNAi plates to develop into gravid adults and were then used for a second TEL on the same RNAi conditions. Progeny from the second-generation TEL were then used for the lifespan assay. RNAi bacteria (HT115) is AMP resistant and can grow slowly on AMP lifespan plates during the assay. To maintain equivalent amounts of bacterial availability over the course of the lifespan, RNAi lifespan plates are seeded with 2x concentrated bacteria for fed conditions and 0.1x concentrated bacteria for DR conditions, from a starting optical density (OD_600_) of 3.0.

Slide Microscopy

Fluorescent images in this study were taken using Leica Application Suite X (LASX) software and a Leica scope with >15 worms per treatment at ≥35x magnification. For representative images displayed in the figures, worms were paralyzed in 0.5M sodium azide (NaN3) and imaged once residual liquid evaporated and the worms clumped together. For quantification of these images and for imaging screens, the same method was used but after the residual sodium azide had evaporated, individual worms were gently pushed apart until they were no longer touching one another. Fluorescent mean comparisons were then quantified using custom R code to identify individual worms and measure fluorescent intensity. The mean fluorescent intensity of the background (all non-worm pixels) was subtracted from the mean fluorescence of each worm. All outputs from this R code were manually quality controlled to eliminate any non-worm objects that were misidentified. This code will be provided upon request to the corresponding author. Data were plotted by R version 4.3.1 and Adobe Illustrator 2022. The brightness of fluorescent images of clumped worms was increased to an equal degree across all images compared to one another in ImageJ bundled with 64-bit Java 1.8.0.

Lawn Occupancy Measurements

For lawn occupancy measurements, worms were grown to gravid adults, then washed off of NGM plates in M9 and pelleted at 1000 *rcf* for 1 minute. At least 10 worms per plate were then pipetted onto the edge of a 35mm diameter NGM plates seeded with 100µL of fed, fed + touch, DR, and DR + touch media (see Fake Lawn Preparation). 5 35mm plates were used for each condition for each replicate. Worms were left on these conditions for 24 hours, and a picture of each plate was taken using Leica Application Suite X (LASX) software and a Leica scope at 7.3x magnification at 1, 3, 6, and 24 hours after placement on conditions. For fed, fed + touch, and DR + touch conditions, the Cell Counter application on image J was used to click on and count each worm in the lawn region and off the lawn region (FIJI Is Just ImageJ bundled with 64-bit Java 1.8.0). For DR conditions, the average area covered by a fed lawn was calculated as a circle with a diameter of 850 pixels. A circle of this size was drawn in ImageJ and placed in the center of the image of the plate. Worms within the circle were counted as “on the lawn” and worms outside of this circle, near the plate perimeter, were counted as “off the lawn”. Data were plotted by R version 4.3.1 and Adobe Illustrator 2022.

Statistical Analysis

Bar plots show data points representing individual worms, with the height of the bar indicating the mean fluorescence in a given condition normalized to the mean of the fed or DR control of that experiment. Different shaped points indicate distinct biological replicates taken across 3 different experiments. The error bars represent SEM centered on the mean. Comparisons between more than two conditions in imaging and lawn occupancy experiments were done using ANOVA. For comparisons between two conditions, a Wilcox rank sum test (two-sided, unpaired) was used with a Bonferroni p-adjustment to account for 3 biological replicates. p values are **p* < 0.05, ***p* < 0.01, ****p< 0.001*, and *****p<*0.0001. For lifespan assays, the statistical groupwise and pairwise comparisons across survivorship curves were performed using the survfit survival analysis package in R. *P* values comparing two survival curves were acquired using the log-rank analysis. *P* values indicating an interaction between multiple experimental variables and across >2 survival curves were performed using the survivalMPL package in R to perform a Cox regression analysis between covariates of interest. Tests done with data from multiple biological variables were adjusted with a Bonferroni p value correction. p values are **p* < 0.05, ***p* < 0.01, ****p< 0.001*, and *****p<*0.0001. Exact *N*s for each experiment are included in the source data files and minimum *N*s for each plot are included in the figure legends.

Data Availability Statement

All raw data and statistical analyses of this data are included in this study and its corresponding supplementary information and source data files. If raw data files are needed in a different format, the corresponding author will provide access to this data upon a reasonable request.

Obtaining Biological Materials

All worm strains generated in this study are available to be shared upon request from the corresponding author.
